# KPB82 potently and selectively inhibits ACER2 and protects tumor and primary cells from doxorubicin-induced cytotoxicity

**DOI:** 10.64898/2026.09.29.754712

**Authors:** Arpita Devi, Ruijuan Xu, Zdzislaw Szulc, Ian Kim, Mohamed Salama, Krishna P. Bhabak, Christoph Arenz, Daniel Canals, Yusuf A. Hannun, Cungui Mao

## Abstract

Alkaline ceramidase 2 (ACER2) catalyzes the hydrolysis of ceramides into sphingosine and free fatty acids, playing crucial roles in regulating various biological processes, including but not limited to cell proliferation, survival, and the DNA damage response. However, a pharmacological tool that regulates its catalytic and signaling functions remains to be developed. In this study, N-[(1S,2R)-1-hydroxy-1-phenylpropan-2-yl]dodecane-1-sulfonamide (KPB82) was identified as a potent and selective ACER2 inhibitor. *In vitro* enzymatic assays showed that KPB82 inhibited ACER2 with an IC_50_ of 190 nM, while displaying minimal activity against acid and neutral ceramidases (IC_50_> 100 µM) and no significant inhibition of alkaline ceramidase 1 or alkaline ceramidase 3. Enzymatic kinetic studies revealed that KPB82 is a competitive inhibitor of ACER2. The three-dimensional structure of ACER2 was modeled in the AlphaFold3 server, revealing key Zn²⁺-coordinating histidine residues and a substrate-binding pocket. Molecular docking studies demonstrated that KPB82 binds ACER2 within this pocket, forming hydrogen bonds with the catalytically important amino residues, Asp93 and His215, and interacting with several hydrophobic residues. Lactate dehydrogenase release assays revealed that KPB82 significantly reduced cytotoxicity in HeLa cells overexpressing ACER2, as well as in HCT116 cells, RT4 cells, and human umbilical vein endothelial cells treated with the DNA-damaging agent doxorubicin. Lipidomic analysis showed that KPB82 inhibited sphingosine production in response to ACER2 overexpression or doxorubicin treatment. Collectively, these findings demonstrate that KPB82 is a potent and selective ACER2 inhibitor capable of suppressing ACER2-dependent biological processes and highlight its potential as a tool for studying ceramide metabolism and ACER2-mediated signaling pathways.

## INTRODUCTION

Ceramidases are a group of enzymes that hydrolyze various ceramide species into sphingosine (SPH) and fatty acids (1). Based on their pH optima for enzymatic activity, ceramidases are classified into acid, neutral, and alkaline ceramidase groups (2). Acid ceramidase (aCDase), encoded by the *ASAH1* gene, has an optimal pH of ∼4.5 and preferentially hydrolyzes unsaturated medium-chain ceramides (3, 4). Neutral ceramidase (nCDase), encoded by *ASAH2*, exhibits optimal activity at pH 7–9 and favors long-chain ceramides as substrates (5). Alkaline ceramidases 1, 2, and 3 are encoded by the *ACER1*, *ACER2*, and *ACER3* genes, respectively, and function optimally at pH 8–9, each with distinct substrate specificities (2). ACER1 preferentially hydrolyzes very long-chain unsaturated ceramides (6), while ACER2 displays broad substrate specificity, acting on both long-chain and very long-chain ceramides (7). ACER3 preferentially hydrolyzes long-chain unsaturated ceramides, dihydroceramides, and phytoceramides (8).

Ceramidases have been implicated in regulating various cellular responses, such as cell proliferation, survival, differentiation, and death, by modulating bioactive sphingolipids, ceramides, SPH, and/orsphingosine-1-phosphate (S1P). Dysregulation of ceramidases has been implicated in various diseases, including cancer. *ASAH1* is overexpressed in triple negative breast cancer (9), glioblastoma (10) and multiple myeloma (11), and its overexpression renders cancer cells resistance to chemotherapy, radiation therapy (12, 13), and targeted therapy (proteosome inhibitor, bortezomib) (11). Zhu *et al.* demonstrate that *ASHA2* is upregulated in tumor tissues and tumor-infiltrating myeloid-derived suppressor cells (MDSCs) from colorectal cancer (CRC) patients and that targeting nCDase with its inhibitor inhibits tumor growth in a mouse model of CRC by inducing ferroptosis in MDSCs (14). However, Sun *et al*. showed that it functions as a tumor suppressor in a mouse model of breast cancer by preventing CD8+ T-cell exhaustion (15). ACER1 is downregulated in tumors from patients with head and neck squamous cell carcinoma (HNSCC) compared to adjacent tissues and its expression levels correlate with tumor immune cell infiltration and overall disease-free survival (16), indicating that ACER1 may function as a tumor suppressor in this cancer type. ACER3 is upregulated in tumors from patients with hepatocellular carcinoma (17) and acute myeloid leukemia (AML) (18) and its upregulation has a tumor-promoting role in both cancers. ACER2 is a p53-regulated enzyme that mediates cell fate during DNA damage: its moderate upregulation promotes cell proliferation and suppresses senescence while its drastic upregulation leads to programmed cell death due to SPH accumulation (19, 20). This duality in the cellular response may contribute to its diverse roles in different cancers, promoting diffuse large B-cell lymphoma (DLBCL) (21) and hepatocellular carcinoma (22), while acting as a tumor suppressor in other contexts. Silencing of ACER2 by miR-140-3p activates AKT/mTOR pathway and promotes prostate cancer cell progression *in vitro* (*23*). ACER2 and sphingomyelin phosphodiesterase acid-like 3B (SMPDL3B) are believed to have functional interplay in ovarian cancer as knockdown of ACER2 in cancer cells with heightened SMPDL3B expression inhibited proliferation, migration and invasion of cancer cells (24). ACER2 is overexpressed in tumor tissue of bladder cancer, where it forms a cold tumor microenvironment (25). ACER2 is also upregulated by TIMELESS circadian regulator in ER-positive breast cancer (26). Upregulation of ACER2 not only contributes to cancer but also to the effectiveness of anticancer therapy (27).

Based on their potential to serve as therapeutics for cancer or other diseases, a number of ceramidase inhibitors have been developed over the last two decades (Supplementary table 1). Most of these inhibitors are specific for acid ceramidase, such as B13 (N-[(1R,2R)-2-hydroxy-1-(hydroxymethyl)-2-(4-nitrophenyl)ethyl]-tetradecanamide), SABRAC (2-bromo-N-[(1S,2R)-2-hydroxy-1-(hydroxymethyl)heptadecyl]-acetamide), SACLAC (2-chloro-N-[(1S,2R)-2-hydroxy-1-(hydroxymethyl)heptadecyl]-acetamide), and 2,4-dioxopyrimidine-1-carboxamide derivatives, while some have dual specificity toward both acid and neutral ceramidases, such as DP24c, KPB70, KPB67, and *N*, *N* –dimethylglycine (DMG)-B13 ester prodrugs (LCL522 and LCL596) (28–30). However, there are only two inhibitors that have been shown to inhibit alkaline ceramidase activity, including D-e-MAPP [(1S,2R)-D-*erythro*-2-(N-Myristoylamino)-1-phenyl-1-propanol)] (31) and N-(3-trifluoromethyl-4-chlorophemyl)-2-methylpentanamide (Compound 2) (32). Therefore, it is an unmet need to develop novel alkaline ceramidase inhibitors.

In this study, we have identified KPB82, a sulfonamide analogue of D-e-MAPP, as a selective and potent inhibitor of ACER2. KPB82 inhibits ACER2 with a low IC_50_ while inhibiting ACER1, ACER3, ASAH1, or ASAH2 with a high IC_50_. We also show that KPB82 at low concentrations could protect the cells from the cytotoxic effects of ACER2 overexpression or DOX by lowering the cellular levels of SPH. *In silico* studies predict that KPB82 occupies the substrate-binding site of ACER2 with its sulfonamide moiety forming hydrogen bonds with catalytically important residues of ACER2, thus inhibiting enzymatic activity.

## MATERIALS AND METHODS

### Reagents

Minimal Essential Medium (MEM), RPMI-1640, and McCoy’s 5A media were purchased from Thermo Fischer Scientific (MA, USA). Dulbecco’s Modified Eagle Media (DMEM), Ca^2+^-free phosphate buffered saline (PBS), fetal bovine serum (FBS), trypsin-EDTA, trypsin neutralizing solution, and antibiotics (penicillin/streptomycin solution, blasticidin, and zeocin) were purchased from Invitrogen Corporation (Carlsbad, CA, USA). Endothelial cell culture medium (ECCM) was purchased from Cell Applications Inc. (San Diego, CA, USA). The sphingolipids and the NBD-conjugated ceramide and sphingosine were purchased from Avanti Polar Lipids (Alabaster, AL, USA). Dox was purchased from Sigma (St. Louis, MO, USA). Thin layers chromatographic plates, silica gel columns, and chemicals used for synthesis of KPB82, and its analogs were purchased from Sigma–Aldrich.

### Synthesis of KPB-82 and its varied chain analogs

KPB82 (LCL-847) and its respective chain analogs (LCL-851-853) were prepared in a one-step N-sulfonation procedure as previously described (30). Briefly, (1*S*, 2*R*)-(+)-norephedrine served as the starting material and was reacted with various sulfonyl chlorides of varying hydrocarbon chain lengths. The resulting crude products were purified via silica gel column chromatography using an eluent system of n-hexane/ethyl acetate/acetone (40:10:0.5, v/v/v). Following chromatography, the compounds were crystallized from n-hexane. Purity was confirmed by thin-layer chromatography (TLC) using n-hexane/ethyl acetate (3:1:0.01, v/v/v). The observed retention factors (*R_f_*) were: LCL-847, 0.71; LCL-851, 0.67; LCL-852, 0.62; and LCL-853, 0.58.

Structural verification was performed using electrospray ionization mass spectrometry (ESI-MS) on a Thermo Fisher Scientific TSQ Quantum Access Max triple quadrupole mass spectrometer coupled with a Vanquish uHPLC system. MS analysis was conducted in ESI positive mode with methanol as the solvent, and the results were shown as follows.

KPB-82 (LCL-847): ESI-MS m/z 401.28 ([M+NH_4_]^+^, 100%) and 366.22 ([ MH-H_2_O]^+^, 92%). Calcd. for C_21_H_37_NO_3_S, m/z 383.25 [M].

LCL-851: ESI-MS m/z 373.22 ([M+NH_4_]^+^, 100%) and 338.16 ([ MH-H_2_O]^+^, 75%). Calcd. for C_19_H_33_NO_3_S, m/z 355.22 [M].

LCL-852: ESI-MS m/z 345.17 ([M+NH_4_]^+^, 100%) and 310.17 ([ MH-H_2_O]^+^, 72%). Calcd. for C_17_H_29_NO_3_S, m/z 327.19 [M].

LCL-853: ESI-MS m/z 317.23 ([M+NH_4_]^+^, 100%) and 282.18 ([ MH-H_2_O]^+^, 86%). Calcd. for C_15_H_25_NO_3_S, m/z 299.16 [M].

### Molecular docking

The three-dimensional (3D) structures of ACER1 and ACER2 were modelled in AlphaFold3 server (alphafoldserver.com), and Zn^2+^ and Ca^2+^ were included in the model as these ions are necessary for the catalytic activity of ACERs. The primary sequences of ACER1 and ACER2 were provided as input along with Zn^2+^ and Ca^2+^ and the 3D structures of the proteins were received as output. The 3D structure of KPB-82 (PubChem CID:71458408)and the two dimensional (2D) structures of D-erythro (*e*)-C_16:0_-ceramide (PubChem CID:5283564), D-*e*-C_18:1_-ceramide (PubChem CID: 5283563), and D-*e*-C_24:1_-ceramide (PubChem CID:5283568) were retrieved from NCBI PubChem Compound. The 3D structures of the ceramides and short-chain analogs of KPB82 were generated on Biovia Discovery Studio 2025 using the sketch molecule tool. Molecular docking was performed using default parameters in AutoDock Vina (33) implemented within the PyRx 0.8 virtual screening tool to generate docked complexes with different binding energies, and the docked complexes with lowest binding energies were analyzed for ligand and protein interactions using Receptor-Ligand interaction tool in Biovia Discovery Studio 2025.

### Cell lines and culture

HCT116, RT4, MCF-7, and RH30 cell lines were purchased from ATCC (American Type Culture Collection) and human umbilical endothelial cells (HUVECs) from Thermo Fisher Scientific. HCT116 cells, RT4 cells, and HUVEC were cultured in DMEM, McCoy’s 5A, and ECCM, respectively; and MCF-7 and RH30 cells were cultured in RPMI-1640 medium. Tetracycline (TET)-regulated ACER1 (HaCER1-TET-ON) and ACER2-expressing (HaCER2-TET-ON) cell lines were generated in our lab previously (34, 35) and were cultured in MEM supplemented with blasticidin (5 μg/ml) and Zeocin (25 μg/ml). To induce overexpression of ACER1 and ACER2 in HaCER1-TET-ON and HaCER2-TET-ON cells, respectively, TET was added to medium when cells reached a 90% confluency. AnMCF7subline (MCF7-ASAH2) overexpressing human nCDase were generated in the lab. The ASAH2 open reading frame was cloned into pENTR4-V5 vector (Addgene #17425) to introduce a V5 tag at the N-terminus. The entry clone was then introduced into pLenti CMV Blast DEST vector (Addgene #17451) by using Gateway™ LR Clonase™ II Enzyme mix (Invitrogen). Lentiviral particles were generated in HEK293T cells as previously described (36). MCF7 cells were infected with lentiviral particles, and growth medium was replaced one day after infection. MCF7-ASAH2 cells were selected with blasticidin (25 μg/ml) for 10 days, and ASAH2 overexpression was confirmed by western blotting using anti-V5 tag antibody (Fig. S1). All media were supplemented with 10% FBS and1% penicillin/streptomycin solution.

### Ceramidase assay

Alkaline ceramidase assays were performed using microsomes and different ceramide substrates. ACER1 was assayed in microsomes (20µgproteins per reaction) isolated from HaCER1-TET-ON cells using NBD-D-e-C_12:0_-ceramide as a substrate at pH 8. ACER2 was assayed in microsomes (20 µg proteins per reaction) isolated from HaCER2-TET-ON cells using D-*e*-C_16:0_-ceramide as a substrate at pH 8. Activity of ACER3 was assayed in microsomes (50 µg proteins per reaction) isolated from RH30 cells expressing high basal levels of ACER3 using D-e-C_18:1_ as a substrate at pH 9.4. Activity assay of acid ceramidase (ASAH1) was performed in microsomes (50 µg proteins per reaction) from RT4 cells using D-e-C_12:0_ ceramide as a substrate at pH 4.5. nCDase activity assay was performed using total cell lysates (50µgproteins per reaction) from MCF7-ASAH2cellsusing D-*e*-NBD-C_12:0_-ceramide as substrate at pH 7.4. The reactions were allowed to continue for one hour at 37°C followed by heat deactivation of the reaction. Total lipids were extracted and the formation of SPH and NBD-decanoic acid was analyzed by LC-MS/MS and Thin Layer Chromatography (TLC), respectively.

### Calculations for IC_50_ of KPB-82

IC_50_ of KPB-82 for inhibition of ACER2 activity was calculated using the software Graphpad Prism. The inhibitor concentrations were placed on the X-axis and enzymatic activity on the Y-axis. Non-linear regression was used to fit the sigmoidal dose-response curve. Following this, IC_50_ was calculated by selecting “log(inhibitor) vs. response–variable slope” option.

### LDH release assay

Lactate dehydrogenase (LDH) release from dying or dead cells was assessed using Cytotoxicity Detection Kit PLUS (Roche Life Science) according to the manufacturer’s instructions.

### RT-PCR assay

HUVECs were treated with different concentration of DOX for 24 hours or with 2µM DOX for different time points. RT4 cells were treated with 1 µM and 2 µM DOX for 24 hours. The cells were harvested after treatment and total RNA was extracted using PureLink RNA mini kit (Invitrogen, Thermo Fischer Scientific, Massachusetts, US) as per manufacturer’s protocol. cDNA was synthesized from the extracted RNA and qPCR was carried out using ACER2 and β-actin primer pairs, respectively. The expression of ACER2 mRNA was normalized to that of β-actin.

### Lipid profiling by LC-MS/MS

Sphingolipids from the enzymatic reactions or cells were analyzed by liquid chromatography tandem mass spectrometry (LC–MS/MS) as described (20). Enzymatic reactions were inactivated by heating at 100°C before 50 µl of an internal sphingolipid (SPL) standard mixture (IS) at a concentration of 1 nmol/ml and 2 ml of the lipid extraction solvent containing ethyl acetate/iso-propanol/water (60/28/12%; v/v/v) were added. Cells were collected and washed with ice-cold 25 mM Tris-HCl buffer (pH 7.4) containing 150 mM NaCl and EDTA before 50 µl of IS and 4 ml of the lipid extraction solvent were added. The IS is a mixture of C_17_SPH, C_17_dihydro-SPH, C_17_SPH-1-phosphate, 13C_16_-ceramide (d13/16), 17C_16_-ceramide (d17:1/16), C_17_-ceramide (d18:1/17:0), and 17C_24:1_-ceramide (d17:1/24:1). After centrifugation, each lipid extract was transferred into a new tube, dried under nitrogen gas, and dissolved in 150 μl of a mobile phase solvent containing methanol with 0.2% formic acid and 1 mM ammonium formate. The mobile phase was injected into Spectra C8SR, 150 × 3.0 mm, 3-μm particle size column integrated in a TSQ Quantum Ultra LC/MS/MS system. The mobile phase for the run was methanol containing 0.2% formic acid and 1.0 mM ammonium formate. The target sphingolipids were identified based on their corresponding peaks and retention times. The levels of sphingolipids from the cells were normalized to total inorganic phosphate (Pi) and expressed as pmoles/µmole Pi.

### Statistical analysis

Data are presented as the mean ± standard deviation (SD) of three independent experiments. Statistical analyses were performed using GraphPad Prism (La Jolla, CA). Differences between groups were evaluated using a two-way ANOVA. A *p*-value < 0.05 was considered statistically significant and denoted by asterisks (*), while *p*-values > 0.05 were considered non-significant (ns).

## RESULTS

### KBP82 inhibits ACER2 catalytic activity *in vitro*

KPB82 has been reported as a low-potency inhibitor of acid and neutral ceramidases, but its activity against alkaline ceramidases is unknown. We assayed ACER1, ACER2, and ACER3 activity across a range of KPB82 concentrations and found that it potently inhibited ACER2 (IC_50_ = 190 nM; Fig. 1A). However, KPB82 showed no activity against ACER1 or ACER3 at concentrations up to 100 µM (Fig. 1B, C). While KPB82 did inhibit nCDase and aCDase (Fig. 1D, E), the IC_50_ values were >100 µM, consistent with Bhabak *et al.* (29). These results demonstrate that KPB82 is a highly selective and potent inhibitor of ACER2.

**Figure 1.**
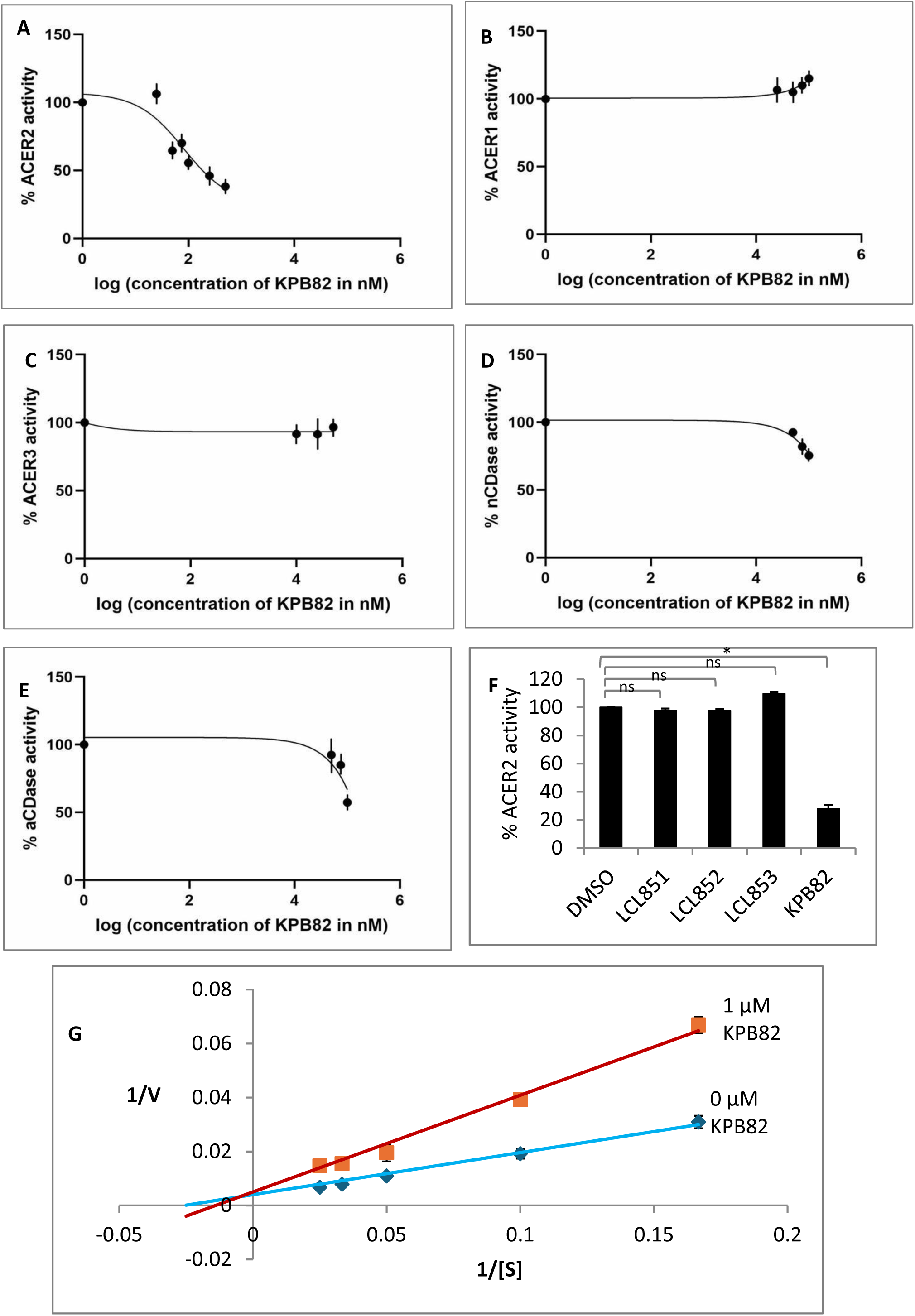
KPB82 potently inhibits ACER2 catalytic activity via competitive or mixed inhibition. (A) Ceramidase activity was measured in microsomes from HaCER2-TET-ON cells overexpressing ACER2 at pH 8.0 in the presence of 50 µM D-*e*-C_16:0_ ceramide and varying concentrations of KPB82; (B) Microsomes from HaCER1-TET-ON cells overexpressing ACER1 were assayed for ceramidase activity at pH 8.0in the presence of 50 µM NBD-D-e-C_12:0_-ceramide and increasing concentrations of KPB82; (C) Microsomes from RH30 cells, which express high basal levels of ACER3, were assessed for ceramidase activity at pH 9.4 in the presence of 50 µM D-*e*-C_18:1_ ceramide and various concentrations of KPB-82; (D) Total cell lysates from MCF7 cells overexpressing ASAH2 were evaluated for neutral ceramidase activity at pH 7.4 in the presence of 25 µM NBD-D-e-C_12:0_-ceramide and different concentrations of KPB82; (E) Total membrane fractions from RT4 cells with high basal levels of ASAH1 were assessed for acid ceramidase activity at pH 4.5 in the presence of 25 µM NBD-D-e-C_12:0_-ceramide and increasing concentrations of KPB82; (F) Ceramidase activity was measured in microsomes from HaCER2-TET-ON cells overexpressing ACER2 at pH 8.0 in the presence of 50 µM D-*e*-C_16:0_ ceramide and 50 µM KPB82, LCL851, LCL852, LCL853, or DMSO (the vehicle); (G) Microsomes from HaCER2-TET-ON cells overexpressing ACER2 were incubated with varying concentrations of D-*e*-C_16:0_ ceramide and 0or 1 µM KPB82 to determine the mode of inhibition. For panels (A–F), control ceramidase activity in the absence of KPB82 was defined as 100%, and activity in the presence of KPB82 was expressed as a percentage of the control. Data represent the mean ± S.D. from three independent experiments performed in triplicate.

To investigate the structural role of the KPB82 alkyl chain, we synthesized and evaluated three short-chain analogues: LCL851, LCL852, and LCL853. These compounds feature chain lengths of 10, 8, and 6 carbons, respectively (Fig. S2). Interestingly, none of the truncated analogues exhibited inhibitory activity against ACER2 at concentrations up to 50 µM (Fig. 1F). These results demonstrate that a specific alkyl chain length is a critical structural requirement for the potent inhibition of ACER2 by KPB82.

To investigate the mechanism of action of KPB82 against ACER2, we performed enzyme kinetic assays. ACER2 activity was measured across a range of substrate concentrations (up to 40 µM) in the presence of 1 µM KPB-82 or the vehicle (control), DMSO, to determine the *K_m_* and *V_max_* values. Our results showed that the *K_m_* increased from 38.8 µM in the control to 72.0 µM in the presence of the inhibitor (Fig. 1G and Table 1). Conversely, the *V_max_* values remained relatively constant at 250.0 and 200.0 pmoles/mg/min, respectively (Fig. 1G and Table 1). These data indicate that KPB82 serves as a competitive inhibitor of ACER2.

### KPB82 occupies a predicted substrate-binding pocket of ACER2 and interacts with catalytical amino acid residues of ACER2

Following our finding that KPB82 is a competitive inhibitor of ACER2, we performed molecular docking to investigate its binding within the catalytic site using the structure predicted by AlphaFold3. ACER2 is characterized by seven α-helical transmembrane domains, consistent with the structure of ACER3 (Fig. 2A). Primary sequence analysis revealed a conservedZn^2+^-binding motif (Fig. 2B), and the AlphaFold3 model predicted that Zn^2+^is coordinated by three histidine residues (His82, His211, and His215) (Fig. 2C).

**Figure 2.**
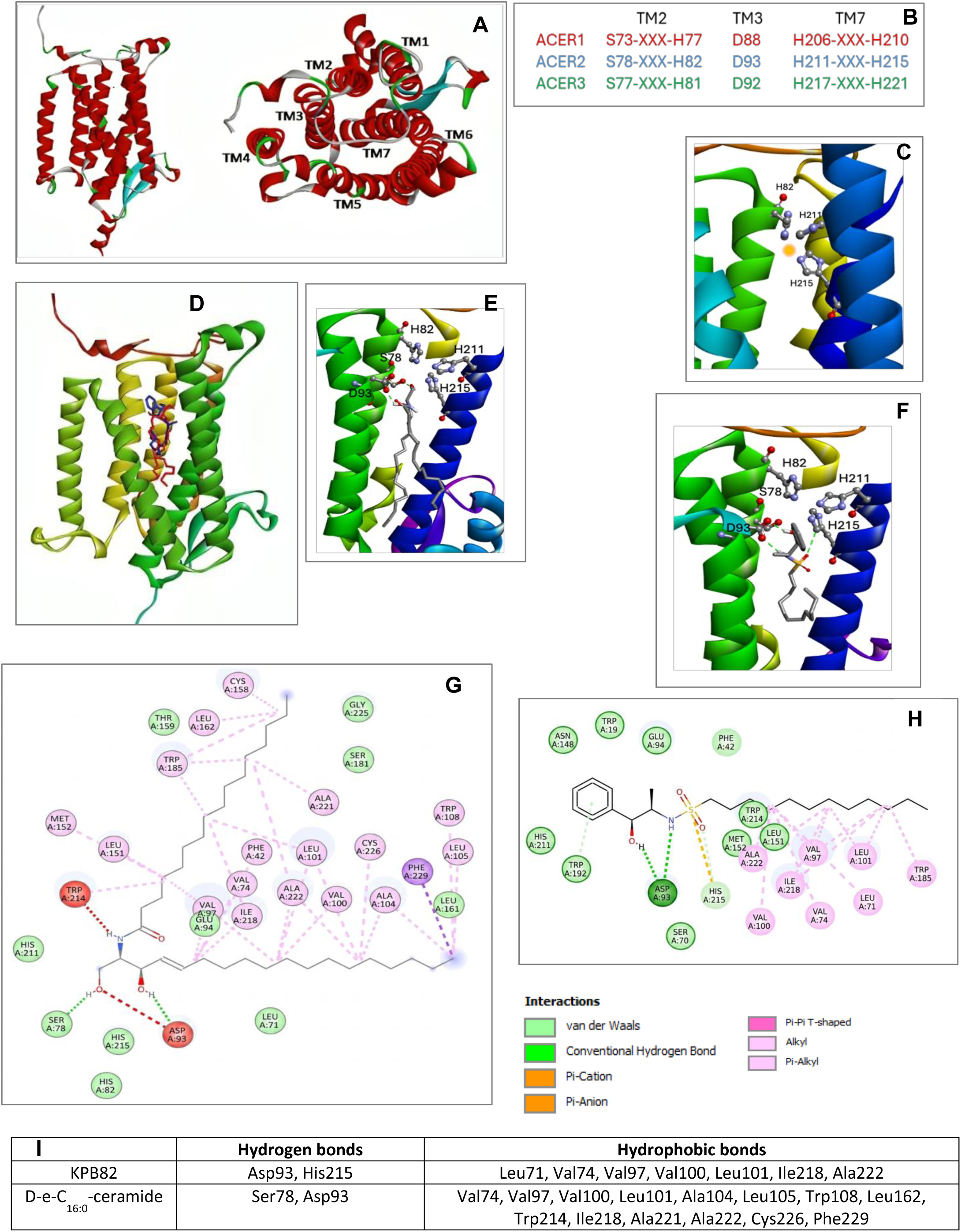
KPB-82 occupies a predicted substrate-binding pocket of ACER2 and interacts with catalytical amino acid residues of ACER2. (A) Side and top views of the AlphaFold3-modeled tertiary structure of ACER2; (B) Sequence alignment showing conservation of Zn²⁺-coordinating residues among ACER1, ACER2, and ACER3; (C) Spatial localization of the three Zn²⁺-coordinating histidine residues in ACER2; (D) Molecular docking shows that both D*-e*-C_16:0_ ceramide (red) and KPB82 (blue) occupy the same binding pocket within ACER2; (E) For D-*e*-C_16_ ceramide, the hydroxyl groups form hydrogen bonds with Asp93 and Ser78 of ACER2; (F) For KPB82, the hydroxyl and amine (–NH) groups form hydrogen bonds with Asp93, and the sulfonyl oxygen forms a hydrogen bond with His215 of ACER2; (G) 2-D representation of ACER2-D-e-C_16:0_ ceramide interaction; (H) 2-D representation of ACER2-KPB82 interaction; (I) KPB82 and D-e-C_16:0_-ceramide form hydrogen bonds with distinct amino acid residues of ACER2 and engage in hydrophobic interactions with several residues.

Molecular docking shows that both KPB82 and D-e-C_16:0_-ceramide, one of the favored substrates of Acer2, sit in the same central cavity surrounded by transmembrane helices TM1, TM2, TM3, TM5, TM6, and TM7 in the 3D model of ACER2 (Fig. 2D). The head groups of both D-*e*-C_16:0_-ceramide and KPB82 situate near the hydrophilic opening of the cavity, where the amide group of D-*e*-C_16:0_-ceramide (Fig. 2E) and sulfonamide group of KPB82 (Fig. 2F) are oriented toward the Zn²⁺ ion. The hydroxyl oxygen of D-*e*-C_16:0_-ceramide forms hydrogen bonds with Ser78 and Asp93 (Fig. 2E, G), and its two hydrocarbon chains sit in the hydrophobic tunnel of the cavity and form hydrophobic interactions with residues Val74, Val97, Val100, Leu101, Ala104, Leu105, Trp108, Leu162, Trp214, Ile218, Ala221, Ala222, Cys226, and Phe229 [Fig. 2G, I]. The oxygen and hydrogen atoms of the sulfonamide group of KPB82 form hydrogen bonds with Asp93, while the sulfonyl oxygen interacts with His215 (Fig. 2F, H). Furthermore, the KPB82 alkyl chain engages in hydrophobic interactions with residues Leu71, Val74, Val97, Val100, Leu101, Ile218, and Ala222 within the same hydrophobic tunnel (Fig. 2H, I). The calculated binding energy is similar for the substrate-enzyme complex (–8.40 kcal/mol) and for the inhibitor-complex (–8.42). These structural interactions, summarized in Fig. 2I, support the prediction that KPB-82 prevents substrate access to the catalytic site through competitive binding.

### KPB-82 occupies the substrate-binding pocket of ACER1 or ACER3 but fails to interact with any catalytic residues

To investigate why KPB82 fails to inhibit the catalytic activity of ACER1 or ACER3, despite ACER2 sharing a high degree of structural similarity with these enzymes, we performed molecular docking of ACER3 and ACER1 with KPB82 and their preferred substrates (D-*e*-C_18:1_-ceramide for ACER3 and D-*e*-C_24:1_-ceramide for ACER1).

In ACER3, D-*e*-C_18:1_-ceramide occupies a central cavity surrounded by the transmembrane helices, TM1, TM2, TM3, TM5, TM6, and TM7 (Fig. 3A). This substrate forms hydrogen bonds with Ser77 and His221 near the hydrophilic opening of the cavity (Fig. 3B). In contrast, KPB82 forms hydrogen bonds with Ser99, Ser228, and Tyr 149 but not any catalytic residue or the Zn^2+^-binding site although it occupies the same cavity (Fig. 3C). Moreover, the hydrophobic interactions between the substrate and enzyme are similar but not identical to those between KPB82 and ACER3. The substrate engages the hydrophobic interactions with residues Val73, Met96, Cys100, Phe103, Val157, Leu179, Phe182, Leu183, Phe186, Trp189, Trp220, and His231 in the hydrophobic tunnel (Fig. 3D) while KPB82 forms hydrophobic interactions with Met43, Val73, Met96, Cys100, Val153, Leu156, Val157, Leu179, and Phe182 (Fig. 3E). For comparison, the interactions of the substrate and inhibitors with ACER3residues are summarized in Fig. 3F. Interestingly, the predicted binding energy for the KPB82–ACER3 interaction (–8.9 kcal/mol) was more favorable than that of the substrate–enzyme interaction (–7.9 kcal/mol).These results suggest that KPB82 fails to inhibit ACER3 likely due to a weak interaction between its hydrophilic head group and the ACER3 substrate pocket, despite favorable binding energy.

**Figure 3.**
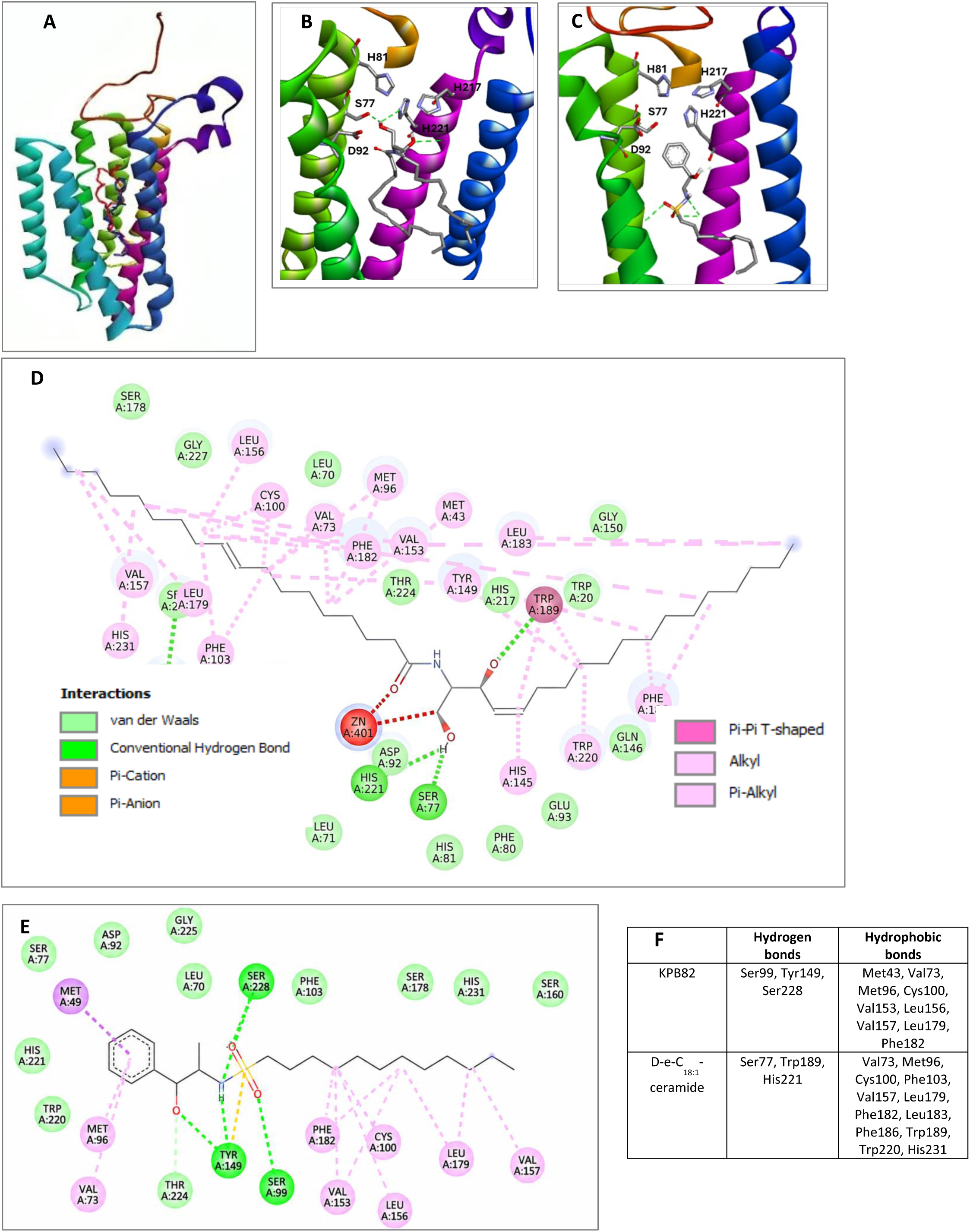
KPB-82 occupies the substrate-binding pocket of ACER3 but fails to interact with any catalytic residues. (A) Molecular docking shows that both D-e-C_18:1_ ceramide (red) and KPB82 (blue) occupy the same binding pocket within ACER3; (B) For D-e-C_18:1_ –ceramide, the head group forms hydrogen bonds with Ser77, His221, and Trp189 of ACER3; (C) For KPB82, the head group forms hydrogen bonds with Ser99. Ser228, and Try149; (D) 2-D representation of ACER3-D-e-C_18:1_-ceramide interaction; (E) 2-D representation of ACER3-KPB82 interaction; (F) KPB-82 and D-*e*-C_18:1_-ceramide form hydrogen bonds with distinct amino acid residues of ACER3 and engage in hydrophobic interactions with several residues.

To investigate the interaction of KPB82 with ACER1, we performed molecular docking analyses using the structure predicted by AlphaFold3.Similarto the structures of ACER2 andACER3, the predicted tertiary structure of ACER2 reveals seven α-helical transmembrane domains (Fig. 4A) and three Zn^2+^-coordinating histidine residues (His77, His206, and His210) (Fig. 4B).

**Figure 4.**
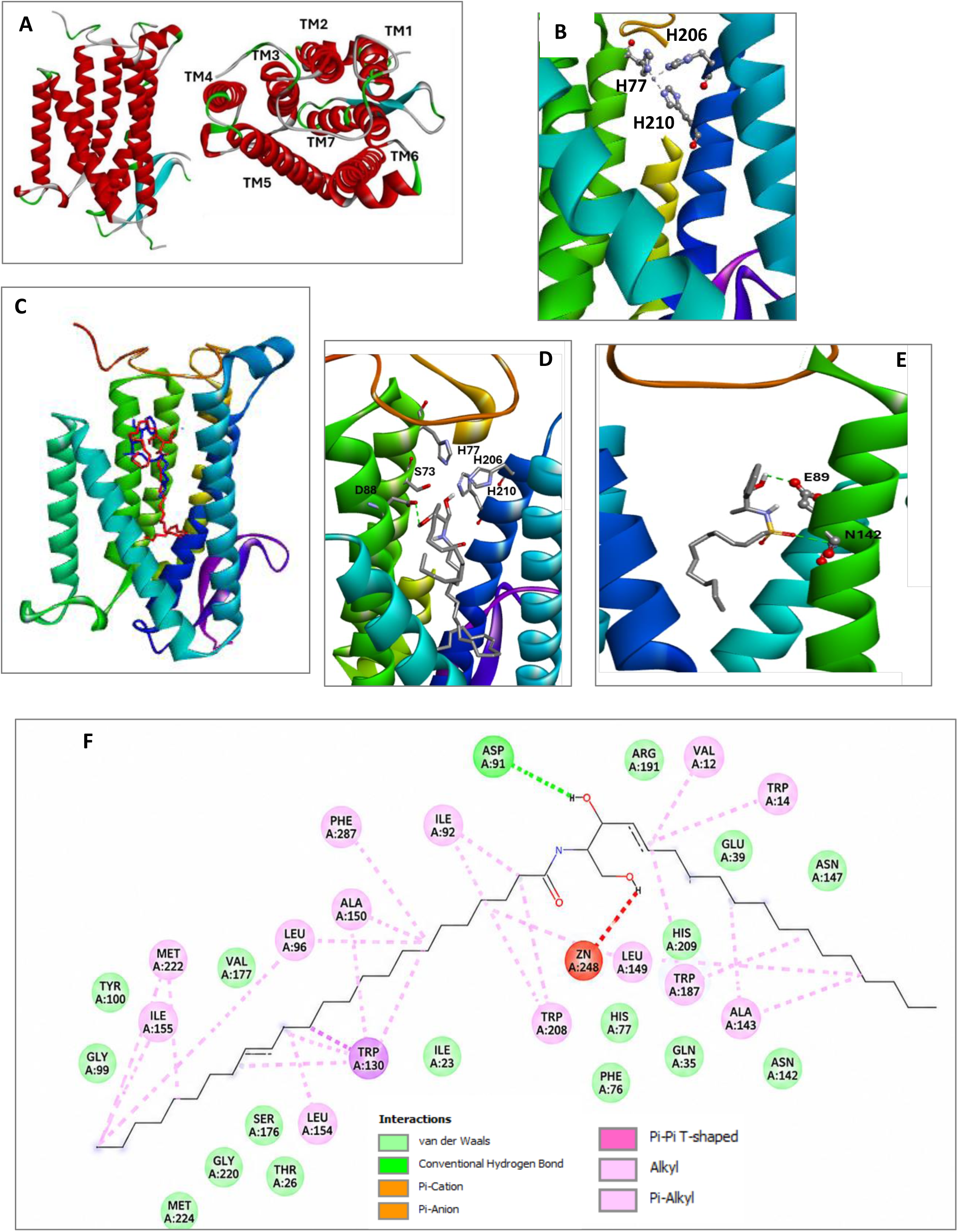
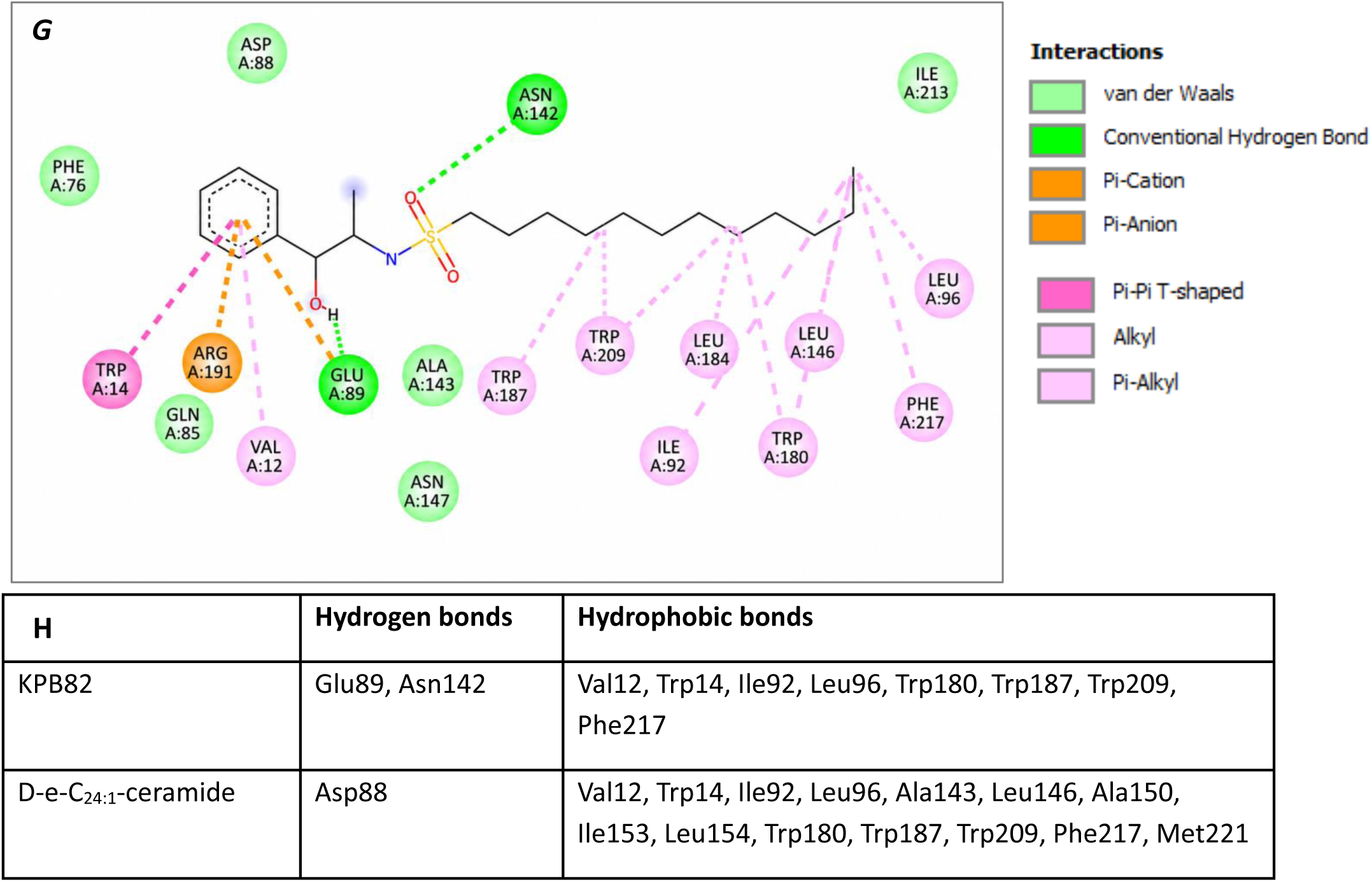
KPB-82 occupies the substrate-binding pocket of ACER1 but fails to interact with any catalytic residues. (A) Side and top views of the AlphaFold3-modeled tertiary structure of ACER1; (B) Spatial localization of the three Zn²⁺-coordinating histidine residues in ACER1; (C) Molecular docking shows that both D-e-C_24:1_ ceramide (red) and KPB82 (blue) occupy the same binding pocket within ACER1; (D) The head group of D-*e*-C_24:1_-ceramide forms the hydrogen bond with Asp88 of ACER1; (E) The head group of KPB-82 forms hydrogen bonds with Glu89 and Asn142; (F) 2-D representation of ACER1-D-*e*-C_24:1_-ceramide interaction; (G) 2-D representation of ACER1-KPB82 interaction; (H) KPB82 and D-*e*-C_24:1_-ceramide form hydrogen bonds with distinct amino acid residues of ACER1 and engage in hydrophobic interactions with several residues.

Docking analysis demonstrated that both KPB82 and D-*e*-C_24:1_-ceramide (a favorite substrate of ACER1) occupy the same central cavity surrounded by the transmembrane helices TM1, TM2, TM3, TM5, TM6, and TM7 in the 3D model of ACER1 (Fig. 4C). D-*e*-C_24:1_-ceramide forms a hydrogen bond with Asp88 near the hydrophilic opening of the cavity (Fig. 4D) and hydrophobic interactions with Val12, Trp14, Ile92, Leu96, Ala143, Leu146, Ala150, Ile153, Leu154, Trp180, Trp187, Trp209, Phe217, and Met221 in the hydrophobic tunnel (Fig. 4F). Interestingly, KPB82 occupies a different position from the substrate within the central cavity (Fig. 4E). Although its N-alkyl chain sits in the hydrophobic tunnel and forms hydrophobic interactions with Val12, Trp14, Ile92, Leu96, Ala143, Leu146, Ala150, Ile153, Leu154, Trp180, Trp187, Trp209, and Phe217 (Fig. 4G, H), its hydroxyl group and sulfonyl oxygen form hydrogen bonds with Glu89 and Asn142, respectively, rather than catalytically critical residues (Fig. 4E, G and H).This binding orientation may explain why KPB82 is predicted to have little or no inhibitory effect on ACER1, even though their calculated binding energy (−6.9 kcal/mol) is the same.

### KPB82 inhibits ACER2-mediated cytotoxicity of doxorubicin by inhibiting SPH generation in tumor and normal cells

Having shown that KPB82 potently inhibited *in vitro* enzymatic activity of ACER2, we tested whether KPB82 can inhibit the role of ACER2 in mediating the generation of SPH in cells. We previously demonstrated that DNA damaging agents, such as DOX, increased expression of ACER2, which in turn increased the levels of SPH in tumor cells expressing wild-type p53 (20). Here we show that treatment of HCT116 cells with DOX increased the cellular levels of SPH as compared to treatment with the vehicle (DMSO), and the increase in SPH levels was reduced significantly in the presence of KPB82 (Fig. 5A). RT4 cells have high basal levels of ACER2 but its expression was not further increased by DOX treatment due to a p53 mutation in these cells (Fig. 5B). However, KPB82 could decrease SPH levels in these cells regardless of DOX treatment (Fig. 5C). Taken together, these results suggest that KPB82 can inhibit the role of ACER2 in mediating the generation of SPH in tumor cells expressing the wild-type p53 or p53-mutant cells expressing high basal levels of ACER2.

**Figure 5.**
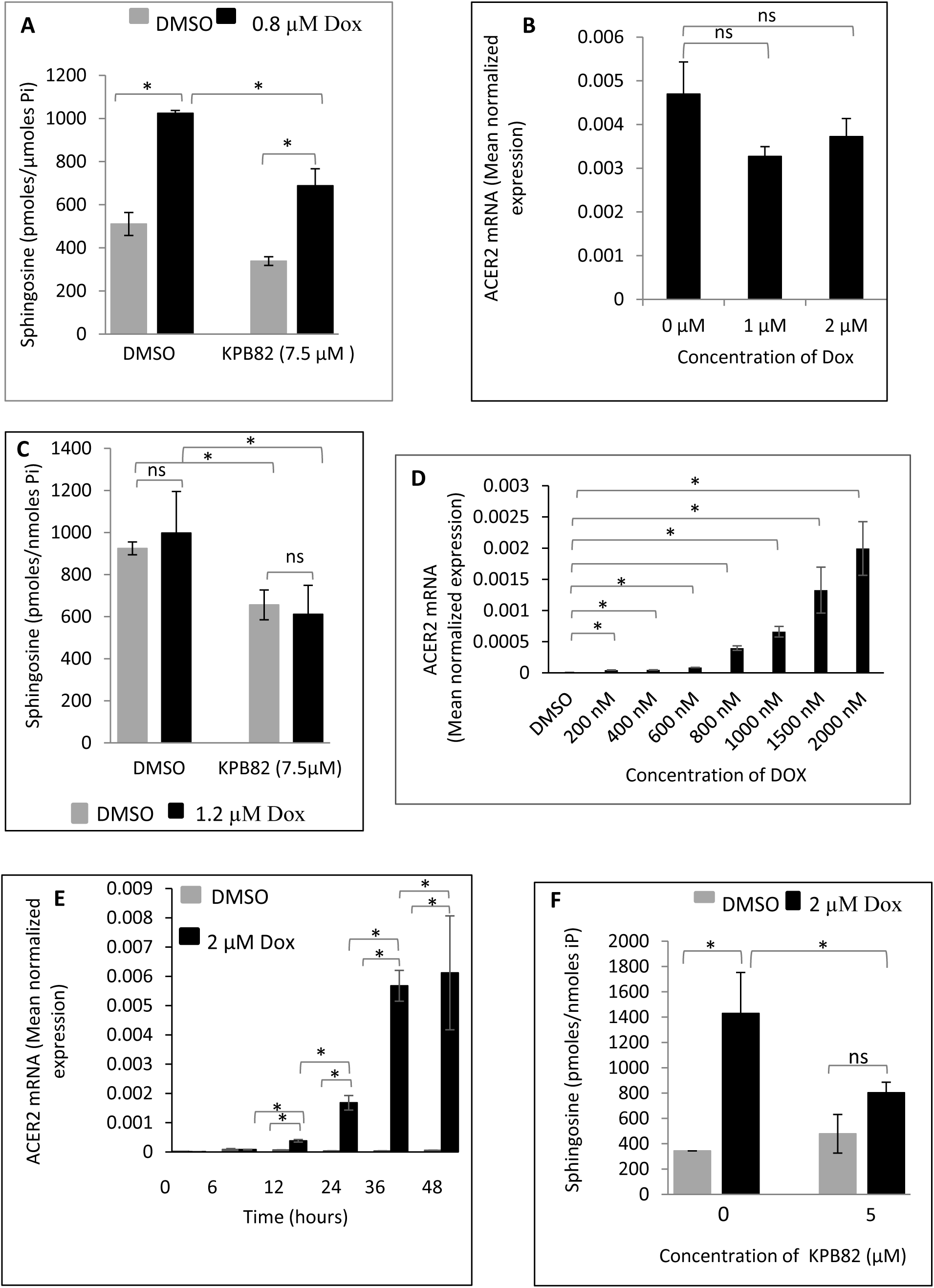
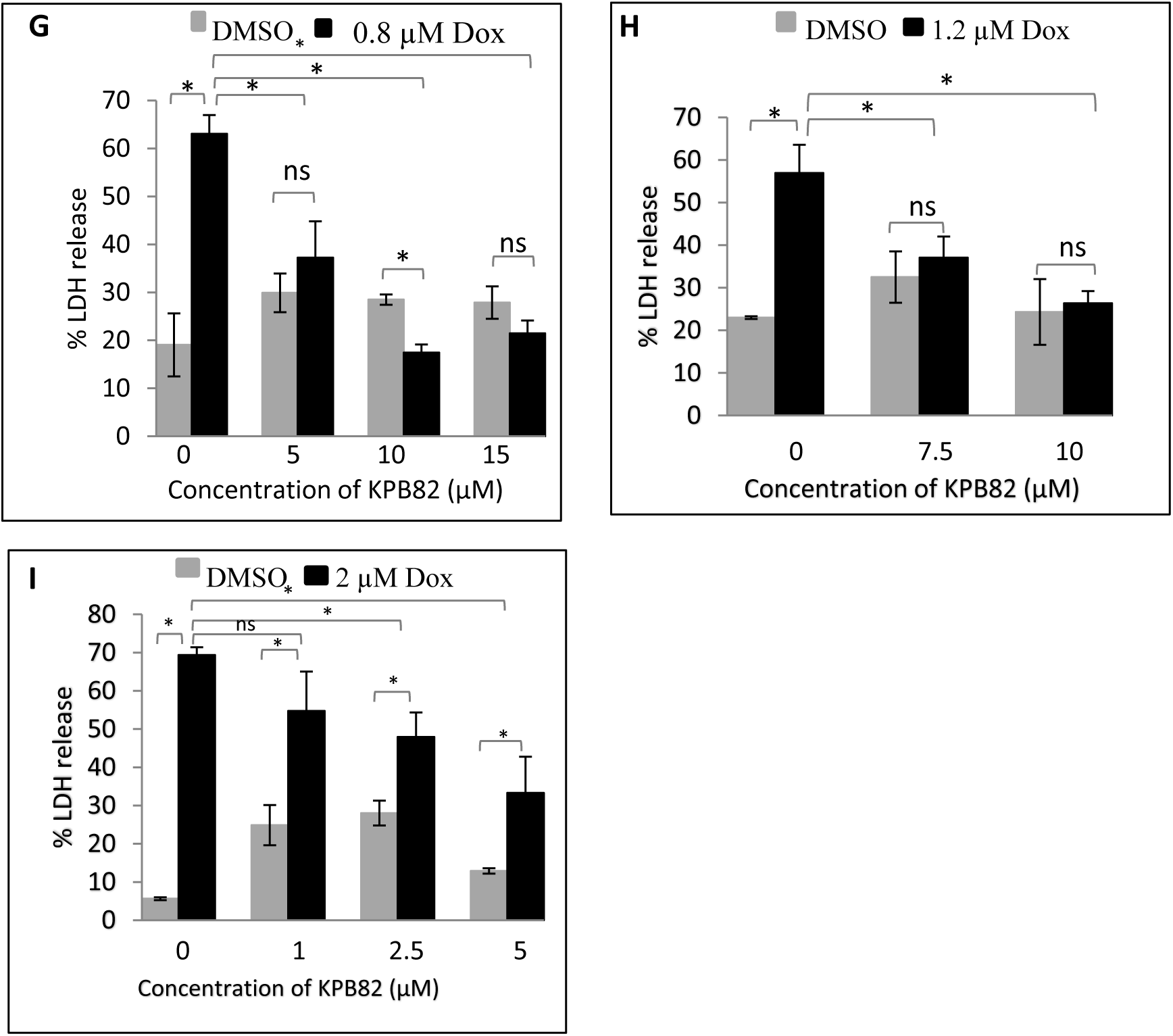
KPB82 inhibits ACER2-mediated cytotoxicity of doxorubicin by inhibiting SPH generation in tumor and normal cells. (A) HCT116 cells treated with saline or 0.8 µM Dox in the presence of DMSO or KPB-82before SPH levels were measured by LC-MS/MS; (B) RT-4 cells treated with saline or different concentrations of DOX before ACER2 mRNA levels were measured by qPCR. ACER2 mRNA levels were normalized against expression of β-actin mRNA levels (Mean normalized expression); (C) RT-4 cells treated with saline or 1.2 µM Doxin the presence of DMSO or KPB-82before SPH levels were measured by LC-MS/MS; (D) HUVECs were treated with 0.2–2 µM Dox for 24 hours, and ACER2 transcript levels were quantified by qPCR; (E) HUVECs were treated with 2 µM Dox for various time periods, and ACER2 transcript levels were measured by qPCR; (F) HUVECs treated with saline or 2 µM Doxin the presence of DMSO or KPB-82 before SPH levels were measured by LC-MS/MS; (G-I) HCT116 cells (G), RT4 cell (H), and HUVEC (I) were treated with DMSO or KPB82 at various concentrations in the presence of DMSO or Dox at the indicated concentrations before cytotoxicity was assessed by measuring lactate dehydrogenase (LDH) release. Data represent the mean ± S.D. from three independent experiments performed in duplicate.

Having shown that KPB82 inhibits ACER2-mediated production of SPH in tumor cells, we tested whether this is also the case in normal cells. Here we demonstrated that DOX increased ACER2 mRNA levels in HUVECs in both dose (Fig. 5D) and time dependent manners (Fig. 5E). DOX also increased SPH levels in HUVEC, and this effect was inhibited by KPB82 treatment (Fig. 5F). These results suggest that KPB82 also inhibits ACER2-mediated production of SPH in normal cells.

Having shown thatKPB82 inhibits the role of ACER2 in the generation of SPH, a potent PCD inducer, in cells, we investigated whether it also inhibits the role of ACER2 in mediating PCD in tumor and normal cells. Our previous studies demonstrated that ACER2 mediates PCD in HCT116 cells in response to DNA damaging agents, such as DOX (20). Here we investigated whether KPB82 could also attenuate DOX-induced cytotoxicity in tumor and normal cells. Treatment of HCT116 cells with 800 nM Dox resulted in 63% LDH release. However, treatment of HCT116 cells with Dox in presence of 5, 10, and 15 µM KPB82 decreased LDH release to 37%, 17%, and 21%, respectively (Fig. 5G). Similarly, treatment of RT4 cells with 1.2 µM Dox resulted in 56.94%LDH release in the absence of KPB82 but 37% and 26% LDH release in the presence of 7.5 µM and 10 µM KPB82, respectively (Fig. 5H).Treatment of HUVECs with 2 µM Dox resulted in 69.36% LDH release but presence of 1.0 M, 2.5 µM and 5.0 µM KPB82 could decrease LDH release to 54%, 47%, and 33%, respectively (Fig. 5I). Taken together, these results suggest that KPB82 also inhibits the role of ACER2 in mediating PCD in response to the chemotherapeutic agent, DOX, in tumor and normal cells.

## DISCUSSION

Although KPB82 was originally designed based on its structural similarity to the known alkaline ceramidase inhibitor D-e-MAPP (30), its specific inhibitory activity against human alkaline ceramidases had not been established. In this study, we identified KPB82 as a potent and selective inhibitor of ACER2 using a combinatorial approach of *in vitro* activity assays and molecular docking. Furthermore, we demonstrated that KPB82 effectively blocks ACER2-mediated SPH generation and subsequent cytotoxicity in both tumor and normal cells treated with the chemotherapeutic agent doxorubicin (DOX).

Given that human ceramidases exhibit distinct substrate specificities and pH optima (1), we systematically characterized the inhibitory profile of KPB82 across the ceramidase family. KPB82 inhibits ACER2 at nanomolar concentrations, whereas it affects acid (ASAH1) and neutral (ASAH2) ceramidases only at micromolar ranges. Most notably, KPB82 failed to inhibit ACER1 or ACER3 at concentrations up to 100 µM, underscoring its high selectivity for the ACER2 isoform.

Kinetic analyses suggest that KPB82 acts as a competitive inhibitor (Fig. 1G). We identified the alkyl chain length as a critical determinant of potency (Fig. 1F), while molecular docking confirmed that KPB82 occupies the ACER2 active site in an orientation mimicking endogenous substrates (Fig. 2D). Structurally, KPB82 retains the ceramide backbone but incorporates a sulfonamide linkage and a rigid 1-phenyl group in place of the sphingosine tail, optimizing its fit within the ACER2 pocket.

The three human alkaline ceramidases (ACER1–3) belong to the CREST superfamily and share a conserved, inverted 7-transmembrane (7TM) helical bundle. While their catalytic cores are nearly superimposable, their substrate-binding pockets possess distinct architectures that dictate ligand specificity (Fig. S3). ACER1 features a deep, highly hydrophobic cavity evolved to stabilize very-long-chain ceramides (e.g., D-e-C_24:1_-ceramide). ACER2 possesses a “wide-mouth” generalist architecture, accommodating a broader range of medium-to-long-chain ceramides. ACER3 contains a narrow, restrictive pocket that excludes bulky saturated chains, only fitting “bent” (unsaturated) C18/C20 chains.

The “wide-mouth” entrance of ACER2 easily accommodates the bulky phenyl group of KPB82, while its pocket depth is perfectly calibrated for the inhibitor’s C12 tail. In contrast, the ACER1 “deep well” likely prevents the C12 chain from reaching the catalytic floor effectively. Furthermore, the rigid aromatic ring of KPB82 is sterically hindered by the narrow ACER3 pocket, precluding entry.

While KPB82 and its natural counterpart, D-e-C_16:0_-ceramide, occupy the same central cavity, the sulfonamide group of KPB82 introduces unique electronic and geometric constraints. Both molecules form hydrogen bonds with the catalytic residue Asp93; however, KPB82 uniquely engages His215, a residue critical for coordinating the active-site Zn^2+^ (Fig. 2F). By interacting with His215, KPB82 may destabilize the enzyme’s structural core. Additionally, the sulfonyl oxygens may displace or mimic the catalytic water molecule coordinated by Zn^2+^, effectively “locking” the enzyme in an inactive state. These structural insights provide a robust roadmap for future ligand optimization.

While DOX remains a cornerstone of cancer therapy, its utility is severely limited by dose-limiting toxicities in normal tissues. Our findings implicate ACER2 as a key mediator of DOX-induced damage in primary endothelial cells (Fig. 5I), consistent with its role as a p53-regulated cell death mediator during the DNA damage response.

Critically, pharmacological inhibition of ACER2 with KPB82 attenuated DOX-induced cytotoxicity in normal cells. This suggests a dual-benefit therapeutic strategy: 1) shielding healthy tissues from chemotherapy-induced adverse events and 2) potentially limiting cancer progression by reducing levels of sphingosine-1-phosphate (S1P), a pro-angiogenic and pro-survival factor regulated by ACER2 activity.

In summary, KPB82 is a potent, selective ACER2 inhibitor that bridges the gap between structural sphingolipid biology and therapeutic application. By modulating the ACER2/SPH/S1P axis, KPB82 offers a promising approach to minimizing the side effects of chemotherapy while simultaneously targeting sphingolipid-mediated oncogenesis.

## ACKNOWLEDGMENTS

We thank the Lipidomics Core Facility at Stony Brook University for performing sphingolipidomic analyses. This work was supported, in whole or in part, by the National Institutes of Health Grants R01CA273263 and P01CA097132.

## Abbreviations

aCDase: acid ceramidase
ACER: alkaline ceramidase
nCDase: neutral ceramidase
Dox: doxorubicin
HUVEC: human umbilical vein endothelial cells
LDH: lactate dehydrogenase

