## Supplemental figures S1-3 for "KPB82 potently and selectively inhibits ACER2 and protects tumor and primary cells from doxorubicin-induced cytotoxicity"

**Supplementary table S1. List of ceramidase inhibitors**

| Compound | $IC_{50}$ value | Target |
|---|---|---|
| C6 Urea Ceramide | NA | Neutral ceramidase |
| D-erythro-MAPP | 1-5 µM | Ceramidase |
| AD-2646 | 40 µM | Ceramidase |
| Ceranib1 | NA | Ceramidase |
| Carmofur | 29 nM | Acid ceramidase |
| Acid Ceramidase-IN-1 | 0.166 µM | Acid ceramidase |
| Acid Ceramidase-IN-2 | NA | Acid ceramidase |
| ARN14974 | 79 nM | Acid ceramidase |
| ARN14988 | 12.8 nM | Acid ceramidase |
| SOBRAC | NA | Acid ceramidase |
| SACLAC | NA | Acid ceramidase |
| W000113414_I13 | 6.6µM | Acid ceramidase |

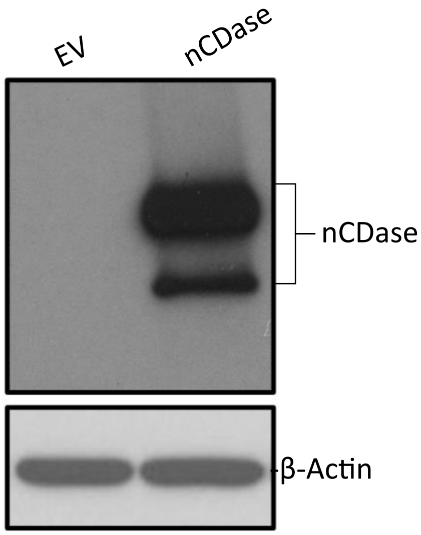


**Figure S1. Neutral ceramidase overexpression in MCF7-ASAH2 cells**

MCF7 cells, stably transfected with an ASAH2-expression construct or empty vector, were cultured as described in the Materials and Methods section. Total cell lysates were prepared from these cells and subjected to western blot analysis using anti-V5 peptide tag antibody.

| Compound ID | 3D representation | 2D representation |
| --- | --- | --- |
| LCL-847 (KPB82) | 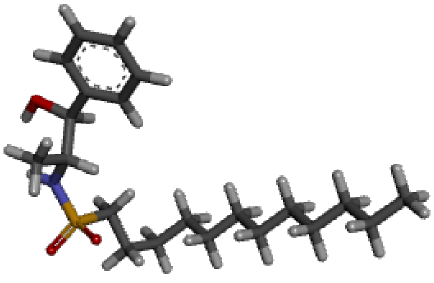 | 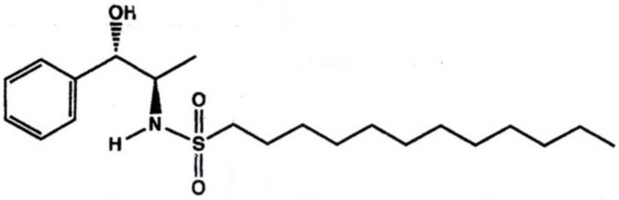 |
| LCL-851 | 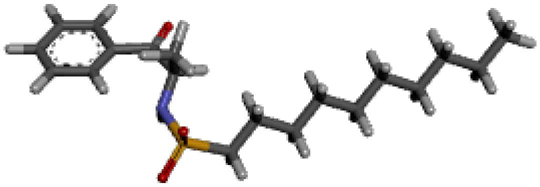 | 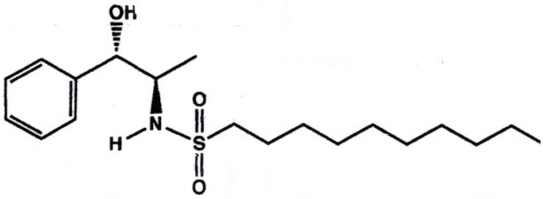 |
| LCL-852 | 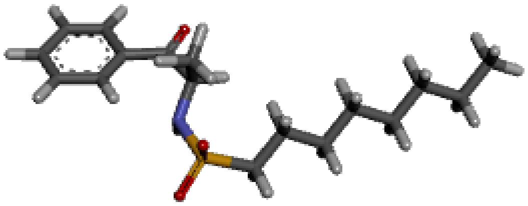 | 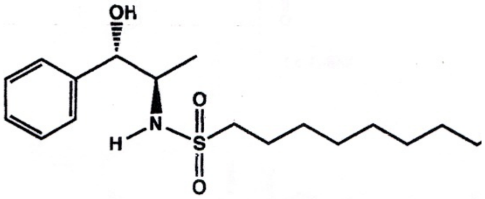 |
| LCL-853 | 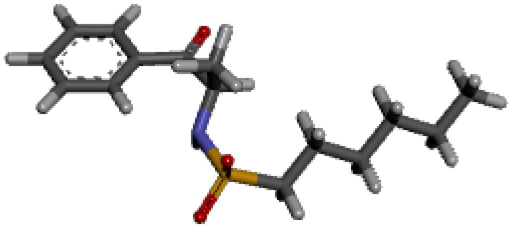 | 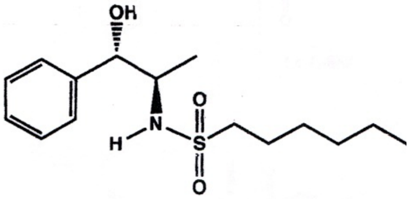 |

**Figure S2. Chemical structures of short-chain analogs of KPB82**

The chemical structures of KPB82 analogs with a 12-carbon (LCL-847), 10-carbon (LCL-851), 8-carbon (LCL-852), and 6-carbon (LCL-853) chains, respectively.

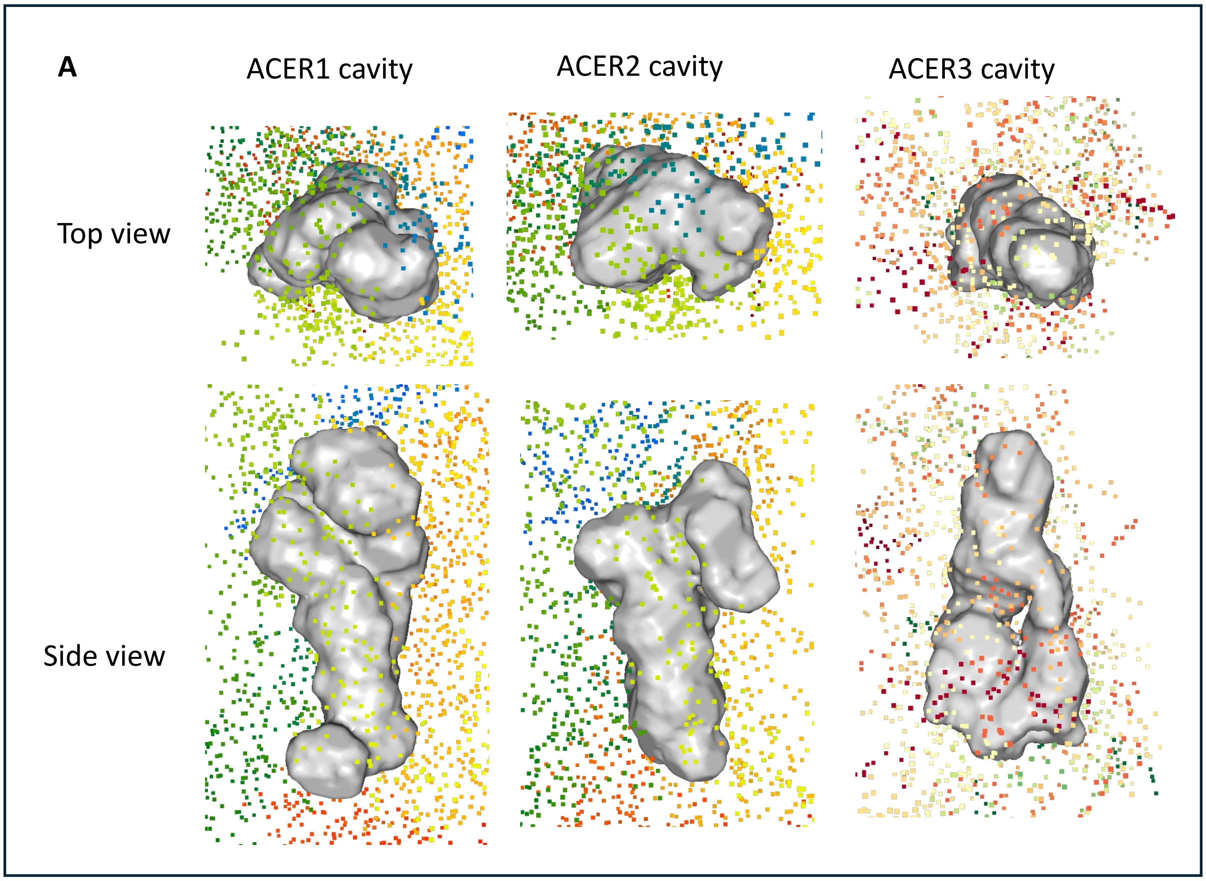


| B | Area (Å²) | Vol. (Å³) | Avg Dep. (Å) | Max Dep. (Å) | Avg Hyd. |
|---|---|---|---|---|---|
| ACER1 | 1011.86 | 1210.46 | 15.88 | 29.01 | -0.72 |
| ACER2 | 762.11 | 1003.54 | 8.35 | 16.94 | -0.65 |
| ACER3 | 615 | 715.18 | 2.61 | 7.85 | -0.38 |

**Figure S3. Distinct architectures of substrate-binding pockets in the alkaline ceramidase family**

The online software KVFinder was employed to predict cavities in the alkaline ceramidases. The largest central hydrophobic cavity from each ACER enzyme was shown (A) and their depths and volumes were listed (B).
